# Crystal structure of β-arrestin 2 in complex with an atypical chemokine receptor phosphopeptide reveals an alternative active conformation

**DOI:** 10.1101/785527

**Authors:** Kyungjin Min, Hye-Jin Yoon, Ji Young Park, Mithu Baidya, Hemlata Dwivedi, Jagannath Maharana, Ka Young Chung, Arun K. Shukla, Hyung Ho Lee

## Abstract

β-arrestins (βarrs) critically regulate signaling and trafficking of G protein-coupled receptors (GPCRs), the largest family of drug targets in the human genome, and there are two isoforms of βarrs: βarr1 and βarr2. Most GPCRs interact with both the heterotrimeric G-proteins and βarrs, inducing distinct downstream signal transduction. However, certain chemokine receptors lack functional G-protein coupling, but they can efficiently recruit βarrs upon agonist-stimulation, and they are referred to as atypical chemokine receptors (ACKRs). Receptor phosphorylation is a key determinant for the binding of βarrs, and understanding the intricate details of receptor-βarr interaction is the next frontier in GPCR structural biology. To date, the high-resolution structures of active βarr1 have been revealed, but the activation mechanism of βarr2 by a phosphorylated GPCR remains elusive. Here, we present a 1.95 Å crystal structure of βarr2 in complex with a phosphopeptide (C7pp) derived from the carboxyl-terminus of ACKR3, also known as CXCR7. The structure of C7pp-bound βarr2 reveals key differences from the previously determined active conformation of βarr1. One of the key differences is that C7pp-bound βarr2 shows a relatively small inter-domain rotation. An antibody-fragment-based conformational sensor and hydrogen/deuterium exchange experiments further corroborate structural features and suggest that the determined structure is an alternative active conformation of βarr2.

## Introduction

G-protein-coupled receptors (GPCRs) are the largest family of receptors on cell membranes and comprise an important class of drug targets. In response to ligand binding, GPCRs activate G proteins as a guanine nucleotide exchange factor, which triggers downstream signaling. To turn off G-protein-mediated GPCR signaling, GPCR kinases phosphorylate the C-terminal tail and/or intracellular loops of GPCRs, which leads to arrestin binding. Although there are over 800 GPCRs in the human genome, only four arrestin genes (arrestins 1-4) have been identified. Among the four arrestin subtypes, arrestin-1 and arrestin-4 are solely related to rhodopsin and cone opsin in the visual system, while arrestin-2 and arrestin-3 (β-arrestin-1 and β-arrestin-2, hereafter βarr1 and βarr2, respectively) are ubiquitously expressed, and they are responsible for interaction with and regulation of nonvisual GPCRs. The interaction of βarrs with phosphorylated receptors transits βarrs to their active state, which leads to desensitization and/or internalization of GPCRs. It is also well established that βarrs critically contribute in a range of downstream signaling responses for many different GPCRs^1^. In addition, βarrs are also recognized as multifunctional and versatile adaptor proteins that bind to and regulate dozens of non-receptor proteins as well^2^.

The interaction of GPCRs and βarrs is typically a two-step process that involves docking of the phosphorylated receptor tail (i.e., the carboxyl-terminus) to the N-domain of βarrs and the interaction of the receptor core (i.e., the intracellular side of the receptor transmembrane bundle) with loops on the convex side of βarrs. While the primary cellular functions of βarrs are broadly conserved across different GPCRs, there is increasing evidence for receptor-specific fine-tuning of βarr functions. Although a clear mechanism for functional diversity of βarrs remains mostly elusive, it has been proposed that different patterns of receptor phosphorylation establish distinct phospho-barcodes on the receptor that fine-tune the interaction pattern and conformational signatures of βarrs, resulting in specific functions^3–6^. To decode how distinct phosphorylation patterns govern βarr conformations and functional outputs, it is essential to visualize the structural details of βarrs in complex with differentially phosphorylated GPCRs or their corresponding phosphopeptides.

There has been significant effort in the recent years to understand the molecular mechanism of βarr activation, including crystal structures of pre-activated arrestin-1^7^, βarr1 in complex with the phosphorylated vasopressin receptor tail (V_2_Rpp)^8^, and a rhodopsin-arrestin-1 fusion protein^9,10^. These structures have revealed major conformational changes upon arrestin-1 and βarr1 activation, such as a significant inter-domain rotation (~20°), disruption of three-element (3E) and polar-core interactions, and reorientation of various loops, including the finger and lariat loops. The V_2_Rpp-bound βarr1 structure also confirmed a previously suggested molecular mechanism, in which the binding of the phosphorylated receptor tail to the N-domain of an arrestin displaces its carboxyl-terminus (Fig. 1a)^11,12^. Furthermore, the crystal structure of a rhodopsin-arrestin-1 fusion protein has also provided structural details for a fully engaged complex, including the interface between arrestin-1 and the receptor core, in addition to a phosphorylation-dependent interaction^9,10^. Single-particle negative-staining-based electron microscopy has also provided direct visualization of the biphasic interaction between the receptor and βarr1 by capturing partially engaged (associated through the receptor tail) and fully engaged (involving the receptor core) complexes^13^.

**Fig. 1.**
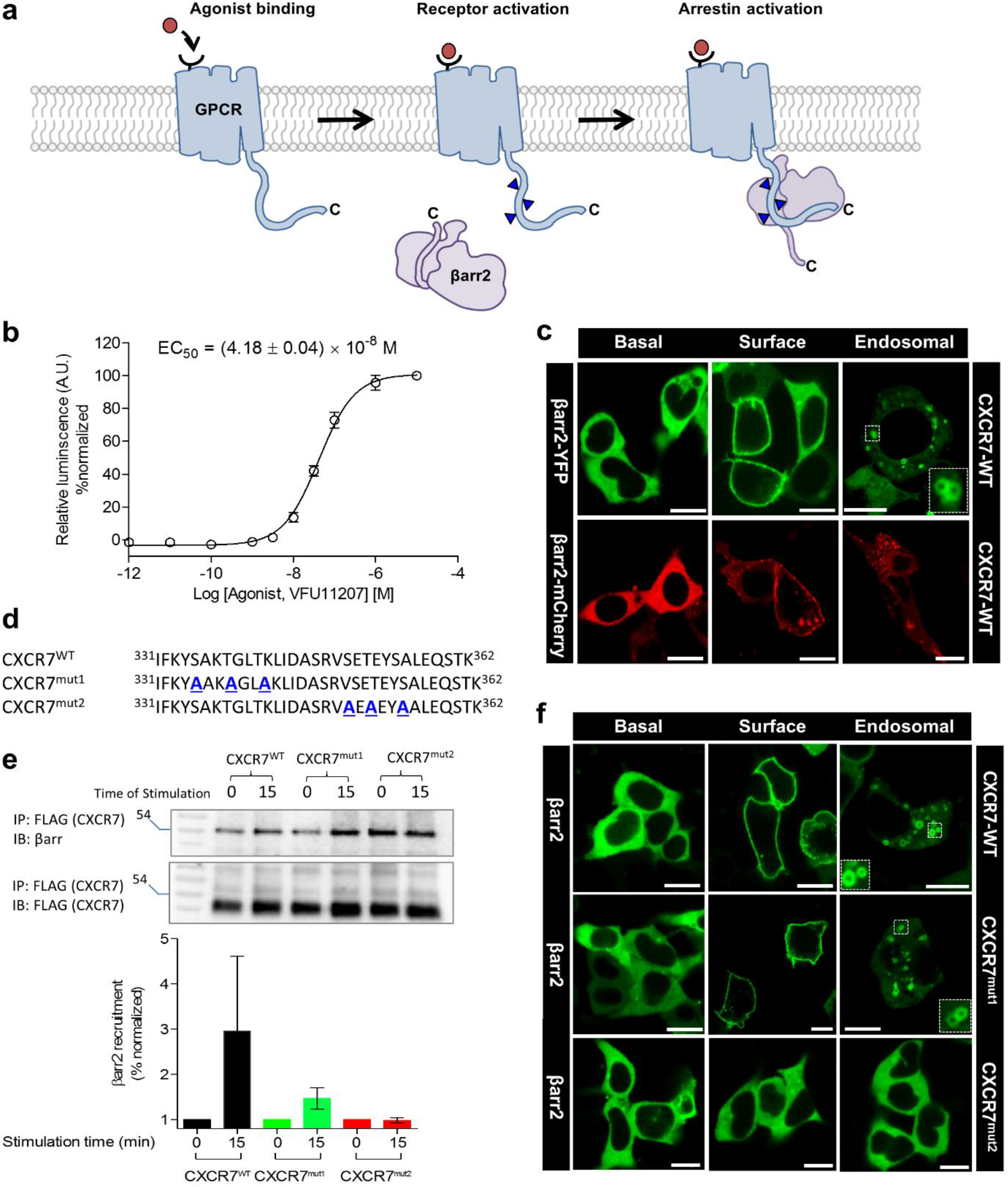
Agonist-induced βarr2 recruitment and trafficking for the human CXCR7. **a**, A schematic representation of the typical βarr2 recruitment to phosphorylated GPCRs. βarr2 interacts with the phosphorylated carboxyl-terminus GPCRs (referred to as receptor tail) first, which displaces the carboxyl-terminal tail of βarr2 docked in the N-domain. Afterwards, βarr2 engages with the intracellular surface of the seven transmembrane bundle (referred to as receptor core) though multiple loops. These two steps of engagement lead to the formation of partially-engaged and fully-engaged complexes, respectively. **b**, Agonist-induced recruitment of βarr2 for CXCR7 as measured by PRESTO-TANGO assay. HTLA cells expressing N-terminal FLAG-tagged CXCR7 were stimulated with an indicated concentration of agonist (VUF11207), followed by measurement of the luminescence output as a readout of βarr2 recruitment. The data were normalized with respect to maximal response (treated as 100%). **c**, Agonist-induced βarr2 trafficking monitored by confocal microscopy. HEK-293 cells expressing CXCR7 together with either βarr2-YFP or βarr2-mCherry were stimulated with a saturating concentration of agonist for the indicated time-points, followed by imaging using the corresponding wavelengths. **d**, Schematic representation of the carboxyl-terminus residues of wild-type CXCR7 and the two phosphorylation site mutants generated in this study. **e**, Co-immunoprecipitation experiments reveal the importance of mutant 2 in βarr2 recruitment. HEK-293 cells expressing CXCR7 constructs and βarr2 were stimulated with a saturating concentration of agonist for the indicated time-points followed by cross-linking using DSP. Subsequently, the receptor was immunoprecipitated using anti-FLAG M1 agarose and the interaction with βarr2 was visualized using Western blotting. The bottom panel shows densitometry-based quantification of data presented in panel **e** normalized with respect to the maximal response for wild-type CXCR7 (treated as 100%). **f**, Confocal microscopy reveals that CXCR7^mut2^ is significantly compromised in inducing βarr2 trafficking upon agonist-stimulation while CXCR7^mut1^ behaves mostly like wild-type CXCR7.

However, activation of βarr2 by the phosphorylated receptor and how it differs from that of βarr1 remain to be structurally visualized. This is particularly important considering that despite ubiquitous expression and high sequence similarity, βarr1 and βarr2 display a significant level of functional divergence^1,14^. For example, some GPCRs bind βarr2 with higher affinity than βarr1 while others recruit both isoforms with similar affinities^15^. Moreover, in some cases, the two isoforms of βarrs also contribute differently toward their conserved functions of receptor desensitization, endocytosis, and signaling^14^. Additionally, for some receptors such as the bradykinin and angiotensin receptors, depletion of βarr2 results in a decrease of agonist-induced ERK1/2 MAP kinase phosphorylation, while depletion of βarr1 enhances it^16,17^. Thus, to fully understand βarr-mediated GPCR regulation and to delineate their functional divergence, visualization of the structural details of βarr2 activation is essential.

Accordingly, in this study we focus on capturing active conformations of βarr2 in complex with phosphopeptides originating from the carboxyl-terminus of a chemokine receptor, CXCR7, also referred to as atypical chemokine receptor 3 (ACKR3). CXCR7, a Class A GPCR, forms a heterodimer with another chemokine receptor, CXCR4, and it has been proposed that it acts as a “scavenger” of its chemokine ligand, CXCL12^18^. It has also been suggested that CXCR7 may represent a natural example of a βarr-biased 7TM receptor because it interacts with βarrs but does not display functional-coupling with heterotrimeric G-proteins^18^. Here, we determine the crystal structure of βarr2 in complex with a CXCR7 phosphopeptide, and the structure reveals key differences with the previously determined structure of V_2_Rpp-bound βarr1. In addition, we also utilize a diverse set of complementary biochemical and biophysical approaches, including site-directed mutagenesis, hydrogen/deuterium exchange mass spectrometry (HDX-MS), and synthetic-antibody-based conformational sensors to extract insights into the activation of βarr2.

## Results and discussion

### Agonist-induced β-arrestin recruitment and trafficking for CXCR7

As mentioned earlier, CXCR7 does not exhibit functional coupling to any of the major sub-types of heterotrimeric G-proteins, although it does efficiently couple to βarr2. We first validated βarr2 coupling and trafficking in HEK-293 cells using a PRESTO-TANGO assay^19^ and confocal microscopy (Fig. 1b-c). We observed that CXCR7 efficiently recruits βarr2 and behaves as a Class B receptor in terms of its trafficking pattern of βarr2 (i.e., receptors are internalized). Based on a recent study that proposed the importance of different phosphorylation codes in GPCRs for βarr binding^10^, we identified two potential phosphorylation-codes in the carboxyl-terminus of CXCR7 with PxxPxxP (pSAKpTGLpT) and PxPxxP (pSEpTEYpS) patterns (Fig. 1d). To identify which phosphorylation code is responsible for βarr2 recruitment to CXCR7, we generated two different CXCR7 phosphorylation-code mutants, referred to as CXCR7^mut1^ and CXCR7^mut2^, that eliminated these two codes sequentially (Fig. 1d). Subsequently, we measured the interaction and trafficking of βarr2 with these two receptor mutants (Fig. 1e-f). We observed that CXCR7^mut1^ behaved essentially similar to a wild-type receptor, while CXCR7^mut2^ exhibited a substantial reduction in βarr2 recruitment and trafficking (Fig. 1f). These findings underscore the relatively important contribution of the second cluster, i.e., the PxPxxP motif, in the βarr2 interaction with CXCR7. A previous study has also demonstrated that the second cluster is more responsible for the internalization and degradation of CXCL12 through CXCR7 than the first cluster^20^.

### Generation and characterization of CXCR7 phosphopeptides

Although the second phosphorylation-code (PxPxxP) was critical for βarr2 recruitment, we synthesized two different phosphopeptides, referred to as C7pp1 and C7pp2, to investigate their interaction with βarr2 and any corresponding structural changes (Fig. 2a). These peptides harbor PxxPxxP and PxPxxP patterns of phosphorylation, respectively (Fig. 2a), and may give us insight into the phospho-code-dependent structural changes of βarr2. These two phosphopeptides exhibited similar binding affinities to βarr2, with dissociation constants (*K*_D_) of 3.08 ± 0.3 μM for C7pp1 and 0.581 ± 0.03 μM for C7pp2, as measured by isothermal titration calorimetry (Fig. 2b). Interestingly however, while C7pp1 displays a monophasic binding with βarr2, the binding parameters of C7pp2 display a better fit using a two-site model, and future studies are necessary to understand the biological significance of this observation, if any.

**Fig. 2.**
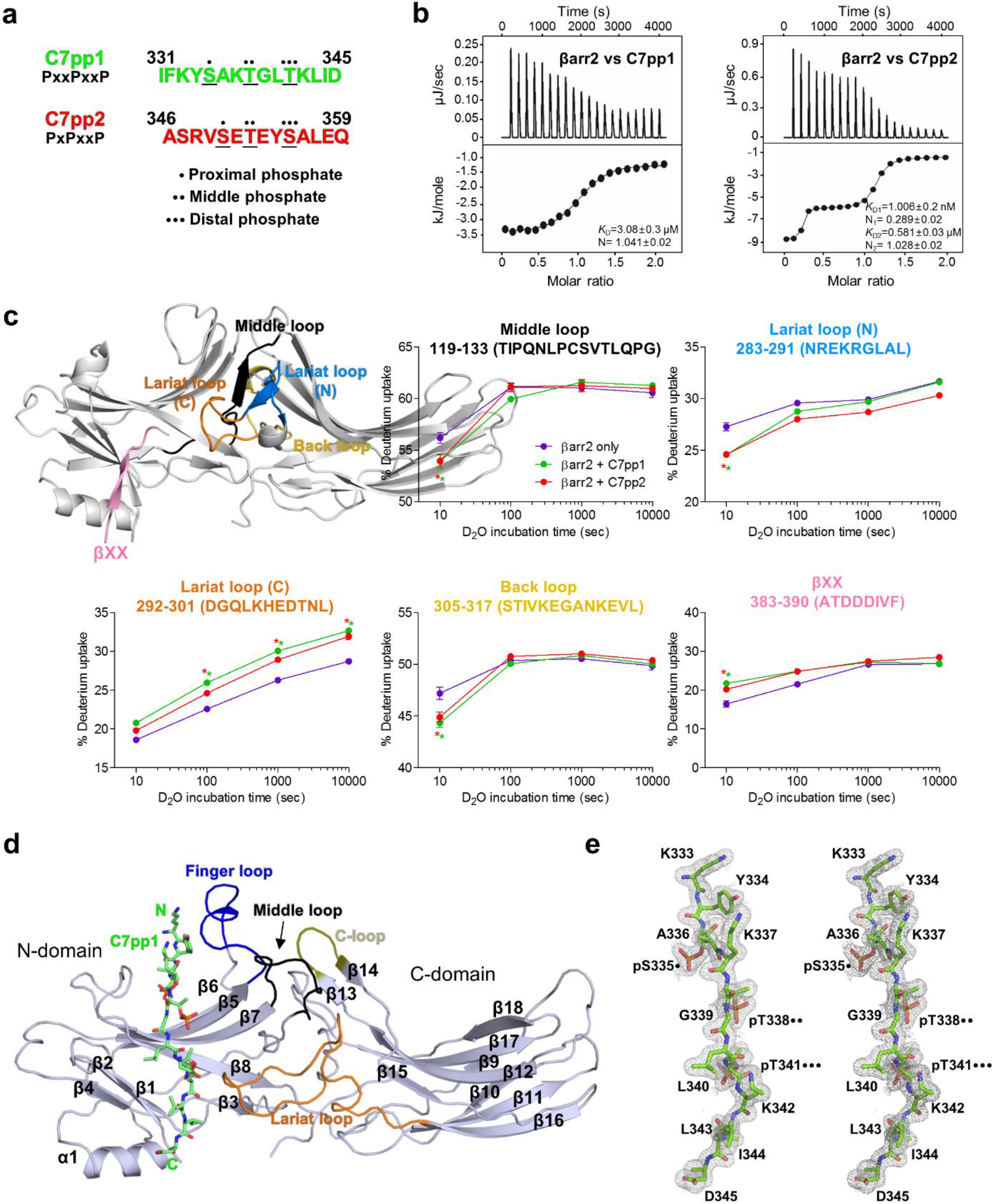
CXCR7 phosphopeptides and crystal structure of βarr2 in complex with C7pp1. **a**, The peptide sequences of the CXCR7 phosphopeptides referred to as C7pp1 and C7pp2 (hereafter, colored in green and red, respectively). The positions of proximal, middle, and distal phosphates of either PxxPxxP or PxPxxP phospho-barcodes are denoted in dots. **b**, Binding affinity of CXCR7 phosphopeptides with βarr2 as measured by isothermal calorimetry. Purified βarr2 was incubated with increasing concentration of the individual phosphopeptides and the binding parameters were calculated based on the dose response curve. The binding constant for each peptide and stoichiometry as observed in three independent experiments is presented here. **c**, HDX-MS profile of βarr2 upon C7pp1 or C7pp2 binding. Regions with altered HDX profile are color-coded on the inactive structure of βarr2 (PDB ID: 3P2D), and the deuterium uptake plots of color-coded regions are provided. Data represent the mean ± standard error of the mean of three independent experiments. The statistical analysis was performed using one-way ANOVA followed by Tukey’s post-test (*p < 0.05 compared to βarr2 alone). Differences smaller than 0.3 Da were not considered significant. **d**, Overall structural snapshot of C7pp1-bound βarr2 high-lighting the loop regions. The C7pp1 peptide is shown as green sticks and the various loops in βarr2, i.e., the finger, middle, lariat, and C-loops in the central crest, are colored in blue, black, orange, and olive, respectively. **e**, The stereo 2Fo-Fc map for C7pp1 is drawn with a 1.0 sigma contour. The positions of proximal, middle, and distal phosphates of the phospho-barcode (PxxPxxP) are denoted in dots.

To understand the structural changes of βarr2 upon C7pp1 or C7pp2 binding, we performed HDX-MS (Fig. 2c). HDX-MS monitored the exchange between the amide hydrogen of a protein and deuterium in the solvent, and the exchange rate was dependent on the solvent exposure and conformational flexibility of the amide hydrogen. The HDX-MS profiles of βarr2 with or without co-incubation of C7pp1 or C7pp2 were analyzed, which showed that C7pp1 and C7pp2 binding induced iconic changes of active arrestins. We observed increased HDX within residues 383-390 containing βXX and residues 292-301 containing the gate loop (the C-terminal part of the lariat loop), implying release of the C-terminus and disruption of the polar core. Additionally, we observed decreased HDX within residues 119-133 containing the middle loop, residues 283-291 containing the N-terminal part of the lariat loop, and residues 305-317 containing the back loop, which may imply the movement of the inter-domain regions. Interestingly, C7pp1 and C7pp2 induced similar HDX changes, suggesting that the overall conformational dynamics between C7pp1- and C7pp2-bound βarr2 were similar although the detailed atomic structures may have differed.

### Crystal structure of phosphopeptide-bound βarr2

To reveal the atomic details of C7pp1- or C7pp2-bound βarr2, we performed X-ray crystallography to obtain high-resolution structures. Although C7pp1 and C7pp2 bind efficiently to the full-length βarr2 (Fig. 2b-c), we used a truncated version of βarr2 lacking the carboxyl-terminal residues 357-410 to facilitate the crystallization of βarr2 in an active conformation. Unfortunately, the crystals of C7pp2-bound βarr2 did not diffract well, whereas we successfully obtained at 1.95 Å resolution the crystal structure of βarr2 in complex with C7pp1 (Fig. 2d and Supplementary Fig. 1a). Therefore, we focus our discussion on the conformational details of C7pp1-bound βarr2.

The crystals of C7pp1-bound βarr2 appeared to be pseudo-merohedrally twinned in the *C*2_1_ space group with a high R_merge_ value, and thus the structure was refined with detwinned data (Supplementary Table 1). The electron density map of residues 331-332 was not observed for C7pp1 (chain U), while nearly all sequences of βarr2 were found to be ordered with the exception of the internal flexible regions (residues 175–181 in chain C and F, respectively) (Supplementary Fig. 1). C7pp1 adopts an elongated loop over the entire length (~35 Å) without severe kinking and is paired with the highly cationic concave surface of the βarr2 N-domain, with a total surface area of 928.4 Å ^2^ buried at the interface (Supplementary Fig. 2).

While interpreting the structural changes in βarr2 upon C7pp1 binding, especially in terms of comparing them with inactive βarr2 or that of V_2_Rpp-bound βarr1, caution is warranted when analyzing the regions involved in crystal contacts that may lead to crystallographic artifacts. Interestingly, the crystallographic asymmetric unit of the βarr2-C7pp1 complex contains six heterodimers of βarr2 and C7pp1, and it shows that the crystallographic contacts of the six molecules are not identical to each other (Supplementary Fig. 1a). Thus, we are able to find at least one solvent-exposed region amongst the six molecules for activation-dependent regions, which allows us to confidently interpret the C7pp1-induced structural changes in βarr2. Moreover, the six βarr2-C7pp1 molecules show essentially similar structures overall when they are superimposed (average root-mean-square deviation of 1.138 Å for the 334 Cα atom pairs) (Supplementary Fig. 1b).

### Smaller inter-domain rotation of C7pp1-βarr2 compared to V_2_Rpp-βarr1

The C7pp1-bound βarr2 structure shares conformational changes similar to other existing active arrestin structures, such as the disruption of the 3E interaction and the polar-core interaction (discussed in Fig. 4b-c). However, the most striking differences between C7pp1-bound βarr2 and other active structures of arrestin-1 or βarr1 is found in the inter-domain rotation angle (Fig. 3a). Pre-activation of arrestin-1 or V_2_Rpp binding to βarr1 induce ~21° and ~20° inter-domain rotation, respectively^7,8^. Even IP_6_−bound βarr2 exhibits ~17° of inter-domain rotation^21^. Like these structural changes, compared to the inactive βarr2 state βarr2 underwent inter-domain rotation in the C7pp1-bound structure (Fig. 3a). However, the inter-domain rotation angle of C7pp1-bound βarr2 is significantly smaller (~8°) than that of βarr1 (~20°) when they are in their final states (Fig. 3a). This data leads us two hypotheses: first, unlike arrestin-1 or βarr1, the receptor-bound βarr2 adopts a structure with smaller inter-domain rotation when it interacts with a phosphorylated receptor C-tail; and second, βarr2 adopts structures with various inter-domain rotations depending on the binding partners.

**Fig. 3.**
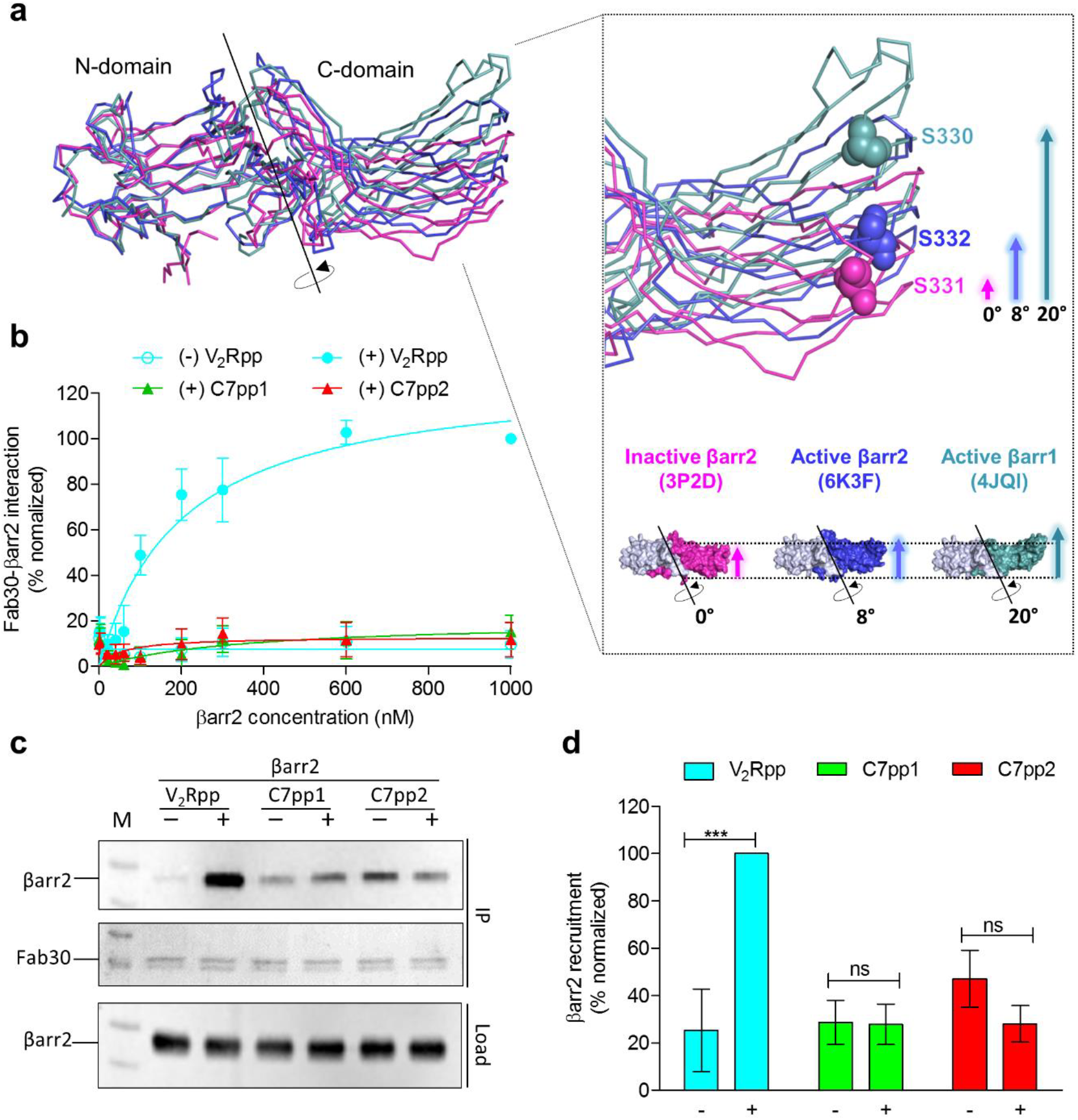
C7pp1-bound βarr2 exhibits a smaller inter-domain rotation compared to V_2_Rpp-βarr1 and the Fab30 sensor corroborates the observation. **a**, Inter-domain rotation comparison of V_2_Rpp-bound βarr1 (light cyan, PDB ID: 4JQI) and C7pp1-bound βarr2 (blue, PDB ID: 6K3F) with the corresponding inactive states of βarr2 (magenta, PDB ID: 3P2D). The N-domains are superimposed, and the rotation axis is indicated in the magnified view of the C-domain. The relative positions of Ser^332^ of active βarr2 (PDB ID: 6K3F) are shown in ball representation as a reference for comparison. The crystal structure of βarr2 in complex with C7pp1 (blue) reveals an inter-domain rotation of about 8**°** compared to the inactive βarr2 structure (magenta), suggesting an intermediate active state. **b**, The Fab30 reactivity pattern corroborates the structural differences between V_2_Rpp-bound βarr1 and C7pp1-bound βarr2. Increasing concentrations of βarr2 in the presence of a saturating concentration of different phosphopeptides were immobilized on an ELISA plate followed by incubation with Fab30 and detection using HRP-coupled Protein L. Data were normalized with the maximal response for V_2_Rpp-βarr2 condition (treated as 100%). **c**, Co-immunoprecipitation experiments further confirm the Fab30 reactivity patterns as observed in ELISA. Purified βarr2 was incubated with a saturating concentration of different phosphopeptides followed by addition of 1.5-fold molar excess of Fab30. Afterwards, Fab30 was immunoprecipitated using Protein-L agarose and the interaction of Fab30 and βarr2 was visualized using Western blotting. A representative image from three independent experiments is shown here. **d**, Densitometry-based quantification of data presented in panel **c** normalized with respect to the maximal response for the V_2_Rpp-βarr2 condition (treated as 100%). Data were analyzed using ONE-WAY ANOVA with Bonferroni post-test (***p<0.001).

**Fig. 4.**
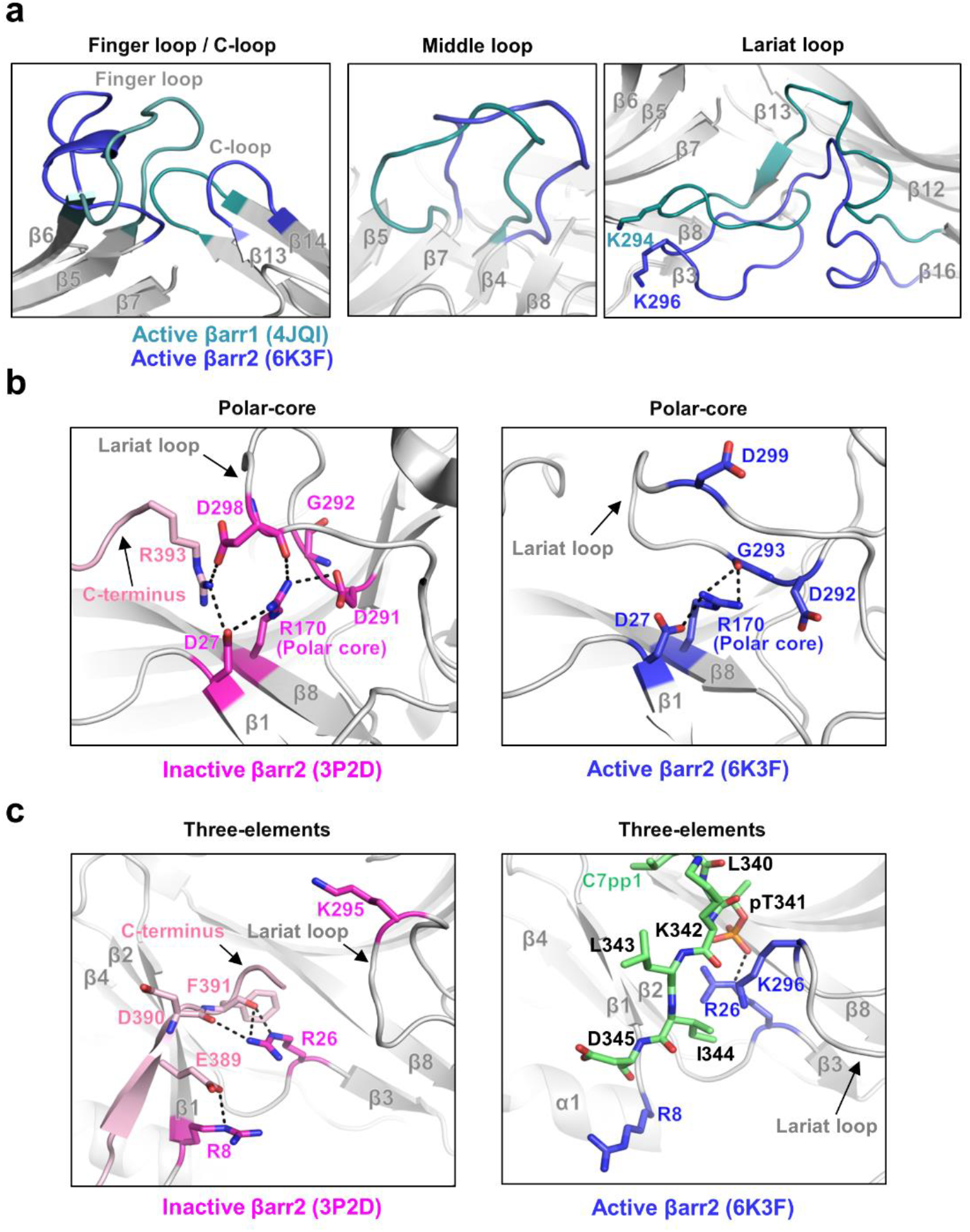
Conformational changes in various loops of βarr2 upon C7pp1 binding as observed in the crystal structure. **a**, Structural comparisons of the finger, middle, lariat, and C loops in C7pp1-bound βarr2 (6K3F) and V_2_Rpp-bound βarr1 (PDB 4JQI). **b-c**, Structural comparisons of polar core and 3E interactions in inactive βarr2 (3P2D, magenta) and C7pp1-bound βarr2 (6K3F, blue), respectively. The C-terminus and C7pp1 are colored in pink and green, respectively.

To test the first hypothesis, we measured the reactivity of a conformationally selective antibody fragment, Fab30, toward C7pp1-, C7pp2-, and V_2_Rpp-bound βarr2. Fab30 efficiently interacted with V_2_Rpp-bound βarr1 and βarr 2, and molecular dynamics simulations have suggested that an inter-domain rotation of more than 15° is most optimal for Fab30 reactivity^22^. We did not observe a significant interaction of Fab30 with C7pp1-bound βarr2 (Fig. 3b-c). This was consistent with the smaller inter-domain rotation observed in the C7pp1-bound crystal structure of βarr2 (Fig. 3a). However, Fab30 interacted robustly with V_2_Rpp-bound βarr2 (Fig. 3b-c). Interestingly, Fab30 also did not interact with C7pp2-bound βarr2 (Fig. 3b-c), which was consistent with the HDX-MS data showing similar conformational dynamics near the inter-domain region between C7pp1- and C7pp2-bound βarr2 (Fig. 2c, residues 119-133, 283-291, 292-301, 305-317, and 383-390). These results suggested that βarr2 adopts different conformations when bound to different R_p_−tails or different activation stimuli, thus rejecting the first hypothesis—the smaller inter-domain rotation in C7pp1-bound βarr2 structure may indicate an inherent propensity specific to βarr2 upon activation.

An alternative hypothesis is that specific phosphorylation patterns, i.e., the number and spatial distribution of phosphates, govern the inter-domain rotation and thereby impart corresponding functional conformation on βarr2. Although such a possibility remains to be explored further, it may explain not only the structural basis of the bar-code hypothesis but also receptor-specific functional outcomes of βarrs. Therefore, we suggest that the current C7pp1-bound βarr2 structure represents an alternative active conformation that may be observed for other receptors as well, depending on the specific phosphorylation pattern. Considering that even partially engaged receptor-βarr conformations are functionally competent, for example, in terms of mediating receptor endocytosis and ERK1/2 MAP kinase activation^21,23^, the current structure has direct implications for understanding the structural details of receptor-βarr interaction, and for ensuring functional responses. It should also be noted that the structure represents the βarr2 conformation in complex with an isolated phosphopeptide without including the core interaction with the receptor. It is also plausible that the core interaction may further fine-tune the conformation of βarr2 including the inter-domain rotation angle.

### Distinct conformational changes of the loop regions in the C7pp1-βarr2 structure

To gain further structural insights into the conformation of βarr2 induced by C7pp1 binding, the active βarr1 structure in complex with V_2_Rpp (4JQI) was compared with the structure of our βarr2-C7pp1 complex (Fig. 4). As discussed above, the N-domain and central loops showed large conformational changes upon activation^24^. The loop regions underwent conformational changes upon C7pp1 binding, and the structures were different in several ways from those of V_2_Rpp-bound βarr1 (Fig. 4a). First, the C7pp1 peptide occluded the inactive conformation of the finger loop lock, promoted outward movement, and induced a helical structure in our crystal structure (Fig. 4a, left panel). This was surprising because the finger loop of the βarr1-V_2_Rpp complex exhibited an extended conformation^8^, and the helical structure of the finger loop was observed when the arrestin was fully docked to the GPCR core. However, it should be noted that HDX-MS analysis did not indicate a helix formation in the finger loop (Fig. 2c), suggesting that the helix observed in the current structure might be a short-lived, very transient state. Second, the middle loop structure was different and did not overlap with those of other arrestins (Fig. 4a, middle panel). Third, the lariat loop moved most closely to the N-domain and made van der Waals contacts with C7pp1 (Fig. 4a, right panel). Lys296 (the corresponding residue of Lys294 in βarr1) belonging to the lariat loop moved toward C7pp1, which might have provided an additional driving force for lariat loop arrangement (Fig. 4a, right panel). Fourth, the C-loop, which was crucial in interacting with GPCR core, resided in similar positions of inactive βarr1 and βarr2, but not the same position (Fig. 4a, left panel). In addition, due to the different inter-domain rotation (Fig. 3), the relative positions of the C-domain were significantly different from each other. We also observed the disruption of the 3E and the polar-core interactions (Fig. 4b-c). Despite the intrinsic flexibility of each loop containing the central crest, the conformations of the six βarr2-C7pp1 molecules in the asymmetric unit matched exceedingly well with each other (Supplementary Fig. 1b), suggesting that each conformation was not derived from a crystallographic artifact, but were the consequences of βarr2 activation by C7pp1 binding. Given that these loops were distributed across the surface of βarr2, different phosphorylation patterns of the GPCR R_p_-tail might induce distinct conformations of βarr2 in a combinatorial manner. Collectively, our structure does not overlap with previously determined structures of βarr1 or βarr2, reflecting the high flexibility of arrestins.

### Distinct binding modes of C7pp1 compared to other R_p_-tails

When we examined the conformations of six C7pp1 peptides in a crystallographic asymmetric unit, two kinds of conformations (chain U vs chains V/W/X/Y/Z) were observed with slightly different modes of βarr2 recognition (Supplementary Fig. 1c). The observation suggested that there could be an ensemble of multiple conformations of C7pp1 when it interacts with positively charged residues distributed on the surface of βarr2 (Supplementary Fig. 2). Given that the N-domain of βarr2 should interact with hundreds of different patterns of the GPCR R_p_-tail, the complex between them might be modular, which has often been observed in disordered proteins^25,26^. The large dependence of electrostatic interactions between βarr2 and R_p_-tails might enable βarr2 to pair promiscuously with hundreds of GPCRs containing differently phosphorylated R_p_-tails.

To investigate how the binding mode of C7pp1 is distinct from those of V_2_Rpp and the rhodopsin C-tail, we compared the conformation of these different structures (Fig. 5a-b and Supplementary Fig. 2c-d). For the structural comparisons, we chose the C7pp1 (chain U) bound to βarr2 with chain A. It was reported that the phosphopeptides overlapped reasonably well when the structure of the rhodopsin-arrestin complex was superimposed with that of the βarr1-V_2_Rpp complex^10^. However, when we superimposed the βarr2-C7pp1 complex with the βarr1-V_2_Rpp complex, the overall conformations of C7pp1 and V_2_Rpp were significantly different (Fig. 5a). The N-terminal part of C7pp1 was closer to the β7/β8 loop compared to that of V_2_Rpp, whereas the C-terminal part of C7pp1 was shorter (Fig. 5a). Especially in the case of βarr1, the N-terminal and C-terminal parts of V_2_Rpp made a continuous β-sheet with β4 and β1, respectively, of βarr1 by anti-parallel stacking. However, those of C7pp1 did not interact directly with either β4 or β1 of βarr2 (Fig. 5a).

**Fig. 5.**
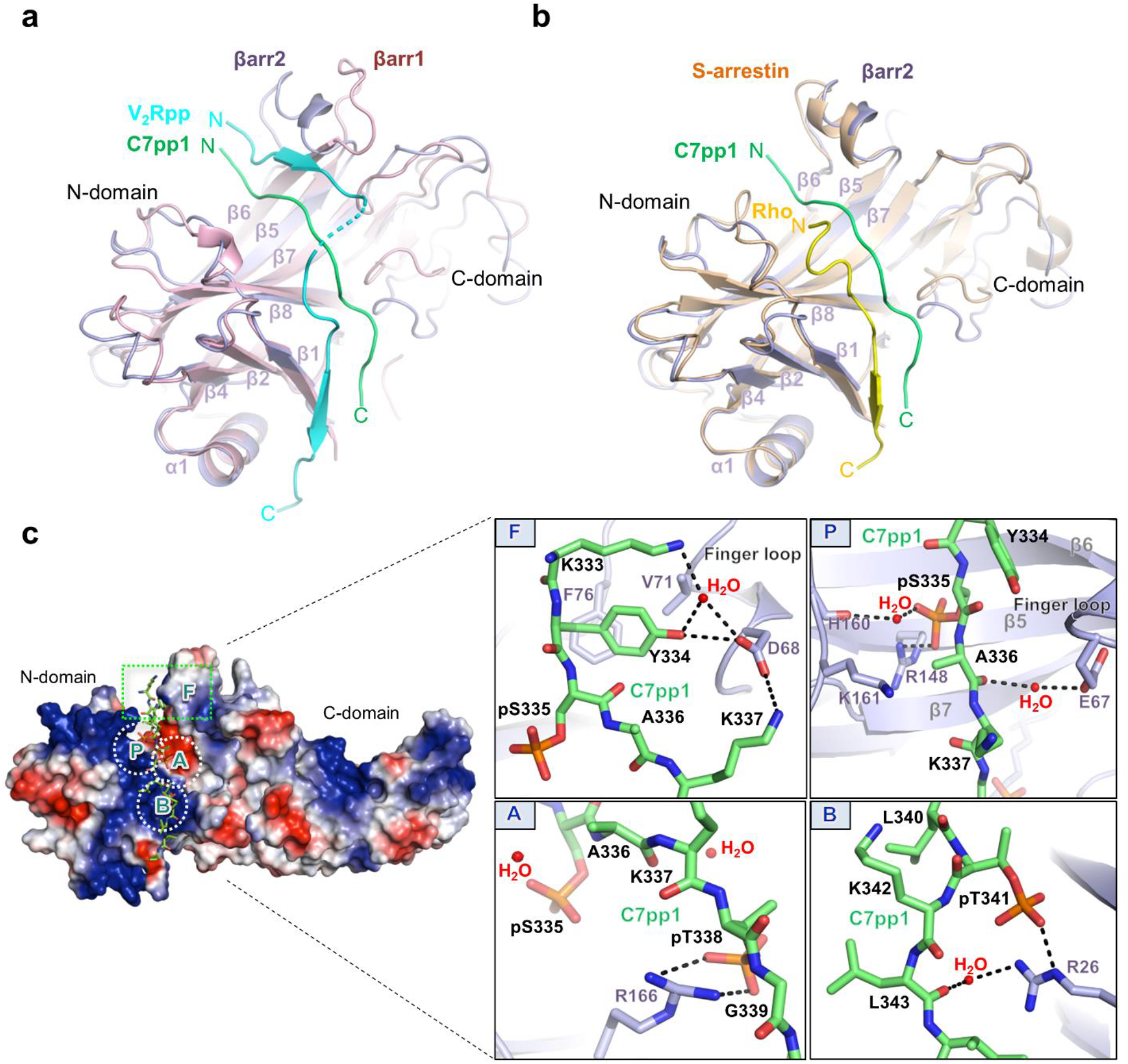
Overall binding mode of C7pp1 to βarr2 with specific interactions of the phosphate groups and activation switches. **a**, An overall distinct binding mode of C7pp1 with βarr2 (6K3F, green) compared to the V_2_Rpp-βarr1 complex (4JQI, light cyan). The N-domains from the crystal structures of the C7pp1-βarr2 complex and V_2_Rpp-βarr1 are superimposed and the respective phosphopeptides are highlighted for comparison. **b**, A comparison of binding modes of C7pp1 with βarr2 (6K3F, green) and the rhodopsin R_p_ tail with S-arrestin (5W0P, yellow) presented in a similar fashion as in panel **a**. **c**, Surface representation for overall electrostatic potential of the C7pp1-bound βarr2 structure. C7pp1 is shown as green sticks. In the positive electrostatic surface of the N-domain, the four hot-spots for C7pp1 binding are shown in the dotted rectangle or circles (F, P, A, and B). The panels on the right represent the detailed interactions at the βarr2-C7pp1 interface and specific interactions of the phosphates with various residues in βarr2.

### Interaction of phosphopeptide with βarr2

Detailed examination of the phosphate-binding sites gave us further insight into the different binding mode of C7pp1 compared to other R_p_ tails. C7pp1 contains three phosphates, which consist of the very frequently observed phosphorylation barcode (PxxPxxP) in the GPCR C-terminus. It has been suggested that three positively charged pockets (pocket A, pocket B, and pocket C) recognize the phosphorylated serine or threonine consisting of the PxxPxxP barcode^10^. In fact, pSer357 and pThr360 (the first and second phosphates) of V_2_Rpp are nearly superimposable with pThr336 and pSer338 of the rhodopsin C-terminal tail, which bind to pocket A and pocket B, respectively^10^ (Supplementary Fig. 3a).

The three phosphates of C7pp1 make extensive contact with positively charged residues on βarr2 (Fig. 5c). The first, second, and third phosphates (pSer335, pThr338, and pThr341) form a salt bridge with βarr2 Arg148 (2.4 Å) (box P), Arg166 (2.8 and 3.4 Å) (box A), and Arg26 (2.5 Å) (box B), respectively. Side chains of many other residues (Lys333, Lys337, Gly339, Lys342, Leu343, and Asp345) except the phosphorylation sites (pSer335, pThr338, pThr341) point in opposite directions from the interface of βarr2 and C7pp1 (Fig. 5c), suggesting that the side chains of other residues do not contribute largely to βarr2 binding.

Interestingly, instead of utilizing the same pockets (A, B, and C) in rhodopsin, the new pocket around Arg148 recognizes the first phosphate (pSer335) (Fig. 5c, box P), while pocket A and pocket B interact with the second and third phosphates (pThr338 and pThr341, respectively) (Fig. 5c, boxes A and B). These results suggest that the binding mode of the PxxPxxP barcode is different in the βarr2-C7pp1 complex. We designated the newly identified pocket, which interacts with the first phosphate (pSer335), as pocket P (Fig. 5c, box P). Next, we checked whether the three pockets (P, A, and B) can accommodate the binding of the PxPxxP barcode, for which they are responsible. It seems that the space between the first and second phosphates can accommodate either one or two residues because the nearby Lys161 (Fig. 5c, box P), which is a strictly conserved residue (Fig. 5c), might interact with the first phosphate of the PxPxxP barcode. It should be noted that the phosphate sensor residues (Arg8, Lys10, Lys11, Lys107, and Lys294 in βarr1) that make contact with the V_2_Rpp phosphates are not involved in the interactions with C7pp1^8,27^ (Supplementary Fig. 3b and 4). The newly identified phosphate binding pocket, pocket P, may be involved in the different conformational changes of C7pp1-bound βarr2 compared to the V_2_Rpp-bound βarr1, which needs further investigation. Together, these data suggest that various GPCR R_p_-tails with different phosphorylation patterns might bind to arrestins differently, which may in turn provide not only the strengths of the interaction but also the ensuing functional outcomes.

### Concluding remarks

As mentioned earlier, an interesting feature of the atypical chemokine receptors (ACKRs) including CXCR7 is their inability to functionally couple G-proteins while maintaining robust interaction with βarrs. Thus, it is tempting to speculate that the conformational differences observed here for βarr2 in complex with C7pp1 when compared to V_2_Rpp-bound βarr1 may reflect a general feature of ACKRs. However, this possibility remains to be experimentally validated in the future for other ACKRs. It is also important to mention that there exists a significant functional divergence between the two isoforms of β-arrestins, βarr1 and βarr2, as mentioned earlier. Thus, it is also plausible that the conformational differences between V_2_Rpp-bound βarr1 and C7pp1-bound βarr2 represent the mechanistic basis of this functional divergence. For example, βarrs have a direct contribution in agonist-induced ERK activation for V_2_R, but for CXCR7, ERK1/2 activation was not observed^18^. Thus, the C7pp1-bound βarr2 structure may represent an alternative conformation that is not competent to activate ERK1/2 but does support receptor endocytosis and thus ligand scavenging, as reported earlier. However, this hypothesis would require additional experimentation in the future, including structure determination with C7pp2 and perhaps with a phosphorylated CXCR7.

In conclusion, we present a C7pp1-bound structure of βarr2 that exhibits key structural differences with the previously determined V_2_Rpp-bound βarr1. These findings shed light on the functional divergence of the two βarr isoforms and also underline the conformational flexibility in βarrs, which allows them to interact with multiple receptors and mediate distinct functional outcomes. Thus, our data may pave the way for developing a better understanding of receptor-βarr interaction and signaling.

## Methods

### Crystallization and data collection

Before crystallization, βarr2_1–356_ (12 mg mL^−1^) in buffer B containing 200 mM NaCl and C7pp1 peptide (70 mg mL^−1^) in 150 mM Tris pH 8.0 were mixed in a 7:1 volume ratio and incubated at 4 °C for 1 h. Crystals of the βarr2_1–356_-C7pp1 complex were grown at 22 °C using sitting-drop vapor diffusion by mixing 1 μL of the protein complex solution with 1 μL of 20% (w/v) PEG 3350, 0.2 M ammonium acetate, and 0.1 M Bis-tris pH 5.5. Crystals were cryoprotected by soaking in Paratone^®^ N oil (Sigma-Aldrich) and flash frozen in liquid nitrogen. X-ray diffraction data were collected at 100 K in 1° oscillations at the BL26B1 beamline of the SPring-8 (Japan). Raw data were processed and scaled using the XDS program suite^28^. Table S1 summarizes the statistics of data collection. The βarr2_1–356_-C7pp1 complex crystal belonged to the space group *C2*_1_, with unit cell parameters of *a* = 91 Å, *b* = 127 Å, and *c* = 206 Å (Table S1).

### Structure determination and refinement

The structure of the βarr2_1–356_-C7pp1 complex was solved by the molecular replacement method using a model of mouse S-arrestin (PDB code 5W0P). A cross-rotational search followed by a translational search was performed using the *Phaser* program^29^. Subsequent manual model building was carried out using the *COOT* program^30^, and restrained refinement was performed using the *REFMAC5* program^31^. Several rounds of model building, simulated annealing, positional refinement, and individual *B*-factor refinement were performed. Table S1 lists the refinement statistics. The asymmetric unit of the βarr2_1–356_-C7pp1 complex contained six molecules of βarr2_1–356_ and peptides, where chains A, B, C, D, E, and F corresponded to βarr2_1–356_, and chains U, V, W, X, Y, and Z corresponded to the C7pp1 peptide. This model included 743 water molecules, and 80.4% of the residues were in the most allowed region of the Ramachandran plot. No electron density was observed for residues 175-181 in chain C and chain F, respectively.

### Accession codes

Crystallographic coordinates of the βarr2-C7pp1 complex have been deposited in the RCSB Protein Data Bank with accession number 6K3F.

## Acknowledgements

The authors thank the staff at Beamline 7A of the Pohang Light Source and BL26B1 beamline of SPring-8 (Japan) for their assistance during the X-ray experiments. This study was supported by a grant from the National Research Foundation of Korea funded by the Korean government (2015R1A5A1008958 and 2018R1A2B2008142) (H.H.L.), (NRF-2019R1A5A202734011) (K.Y.C.), (2018R1D1A1B07040808) (H.J.Y.), and a grant from the Korea CCS R&D Center (KCRC; 2014M1A8A1049296) (H.H.L.). The research program in Dr. Shukla’s laboratory is supported by an Intermediate Fellowship of the Wellcome Trust/DBT India Alliance Fellowship (grant number IA/I/14/1/501285) awarded to AKS, the Science and Engineering Research Board (SERB) (EMR/2017/003804), the Innovative Young Biotechnologist Award from the Department of Biotechnology (DBT) (BT/08/IYBA/2014-3), and the Indian Institute of Technology, Kanpur. Dr. Shukla is an Intermediate Fellow of Welcome Trust/DBT India Alliance, EMBO Young Investigator, and Joy Gill Chair Professor.

## Author contributions

KM and HJY solved the crystal structure. MB designed and generated CXCR7 phospho-site mutations and performed confocal microscopy. MB and HM carried out the co-IP experiments with CXCR7. HM and JM measured the reactivity of Fab30 with C7pp1/2-bound βarr2 using co-IP and ELISA. AKS supervised the experiments performed by MB, HM, and JM, and contributed in writing and editing the manuscript. JYP generated the βarr2 constructs and performed HDX-MS. KYC supervised the experiments performed by JYP and contributed in writing and editing the manuscript. KM, HJY, KYC, AKS, and HHL analyzed the data and wrote the manuscript. HHL directed the teams and all authors edited the manuscript.

## Conflict of interest

The authors declare no competing financial interests.

