## Supporting information for "Crystal structure of β-arrestin 2 in complex with an atypical chemokine receptor phosphopeptide reveals an alternative active conformation"

### This PDF file includes:

Supplementary text
Supplementary Figures 1 to 4
Supplementary Table 1
References for SI reference citations

### Supporting Information

**Cloning, protein expression, and purification.** CXCR7 phosphopeptides (C7pp1 and C7pp2) for crystallization, HDX-MS, and ITC experiments were obtained from NovoPep (Fig. 1a). The *Rattus norvegicus* wild-type βarr2$_{1–410}$ and C-terminal truncated βarr2$_{1–356}$ were inserted into expression vector pET28a. The plasmids were transformed into *Escherichia coli* BL21(DE3)pLysS cells (Invitrogen), and cells harboring the plasmids were grown at 37°C until the optical density (at 600 nm) reached 0.7–1.0 in Luria Bertani (LB) broth containing 70 μg mL$^{-1}$ chloramphenicol and 30 μg mL$^{-1}$ kanamycin. For the crystallization experiment, cells harboring the βarr2$_{1–356}$ were grown at 37°C until the optical density (at 600 nm) reached 1.0 in M9 minimal salt media (Sigma Aldrich) containing 70 μg mL$^{-1}$ chloramphenicol and 30 μg mL$^{-1}$ kanamycin. Further, 0.1 mM isopropyl-β-D-1-thiogalactopyranoside (IPTG) was used to induce protein expression in the cells, after which the cells were incubated for 16 hours at 16°C. Summaries of constructs, primers, restriction enzyme sites, and templates used in this study are included in Table S1. All constructs were verified by DNA sequencing.

For isolation of βarr2$_{1–356}$ protein fused to an N-terminal His$_6$ tag, cells were harvested by centrifugation at 5,000 rpm at 4°C for 10 minutes, and the pellet was resuspended in ice-cold buffer A (20 mM Tris-HCl pH 8.0 and 500 mM NaCl) containing 1 mM phenylmethanesulfonylfluoride (PMSF). The cells were lysed using a microfluidizer (Microfluidics, Westwood, MA, USA), and the lysed cells were centrifuged at 15,000 rpm (Vision V506CA rotor) at 4°C for 30 minutes to separate the supernatant and cell debris. The supernatant was applied to a Ni-sepharose affinity column (GE Healthcare, Little Chalfont, UK) pre-equilibrated with buffer A. Initially, the column was washed extensively with buffer A, after which the protein was eluted using buffer A containing a gradient of imidazole concentrations from 100 mM to 1 M. The eluates were desalted into buffer B (20 mM Tris-HCl pH 8.0 and 5 mM β-mercaptoethanol) containing 100 mM NaCl by using desalting column (GE Healthcare) and further purified by anion-exchange chromatography with Hitrap Q sepharose column (GE Healthcare). The proteins were eluted using buffer B containing 500 mM NaCl in Q column. Further purification was performed by gel filtration on a HiLoad 16/60 Superdex 200 prep-grade column (GE Healthcare), which was equilibrated with buffer B containing 200 mM NaCl. For the $\beta arr2_{1–410}$, the purification steps are the same as $\beta arr2_{1–356}$ construct until application to desalting column (GE Healthcare). After desalting into buffer B, protein was applied to HiTrap heparin column (GE Healthcare) and eluted using buffer B containing 1 M NaCl. Further purification was performed by gel filtration on a HiLoad 16/60 Superdex 200 prep-grade column (GE Healthcare), which was equilibrated with buffer B containing 200 mM NaCl. The homogeneity of the all purified protein was assessed by polyacrylamide gel electrophoresis in the presence of 0.1% (w/v) sodium dodecyl sulfate. The protein solution was concentrated to about 12 mg $mL^{-1}$ using a centricon centrifugal filter unit (Sartorius Stedim). The protein concentration was estimated by measuring the absorbance at 280 nm.

**Isothermal titration calorimetry (ITC).** ITC experiments were performed using Affinity ITC instruments (TA Instruments, New Castle, DE, USA) at 298 K. 100 μM of $\beta arr2_{1–410}$ WT, which was prepared in a buffer containing 20 mM Tris-HCl pH 8.0 and 200 mM NaCl was degassed at 295 K prior to measurements. Using a micro-syringe, 2.5 μL of 750 μM C7pp1 and C7pp2 peptide solutions was added at intervals of 200 s to the $\beta arr2_{1–410}$ WT solution, in the cell with gentle stirring. 30 μM of βarr2$_{1–410}$ WT was prepared in a buffer containing 20 mM Tris-HCl pH 8.0 and 200 mM NaCl was degassed at 295 K prior to measurements.

**ELISA**. For dose response ELISA, purified βarr2 was incubated with 10 fold molar excess $V_2$Rpp/C7pp1/C7pp2 for 30 minutes. Thereafter, varying concentration of peptide bound or unbound βarr2 was immobilized in 96 well MaxiSorp polystyrene plates (Nunc) at room temperature for 1h. The potential non-specific binding sites in the wells were then blocked by incubation with 1% BSA at room temperature for 1h. Subsequently, purified Fab 30 (1 μg/100 μl/well) was added to the wells and incubated at room temperature for 1 h. Wells were washed extensively using 20 mM Hepes pH 7.4, 150 mM NaCl, 0.01% MNG and then incubated with 1:2,000 dilution of HRP coupled Protein L antibody (GenScript). After 1 hour incubation, wells were thoroughly washed and the entire residual buffer removed by blotting on absorbent paper. Thereafter, 3,3',5,5' -tetramethylbenzidine (TMB) (Genscript) substrate was added to each well. Colorimetric reaction was stopped by adding 1M $H_2SO_4$ and absorbance was measured at 450 nm using a Victor X4 plate reader (Perkin-Elmer). All the ELISA data are normalized with respect to the signal for highest concentration of βarr2$^{+V2Rpp}$-Fab30 complex which is treated as 100%.

**Co-immunoprecipitation.** Purified βarr2 (5 μg per 100 μl reaction in 20 mM HEPES pH 7.4, 150 mM NaCl buffer) were activated with $V_2$Rpp, C7pp1, or C7pp2 for 30 minutes at room temperature (25°C) in tumbling conditions. Subsequently, Fab30 (2.5 μg) was added and allowed for binding in room temperature for 1hour followed by addition of pre-washed and equilibrated (in 20 mM HEPES pH 7.4, 150 mM NaCl buffer) Protein L beads to the reaction mixture and additional tumbling at room temperature for 1h. Afterwards, beads were washed 4-5 times with 20 mM HEPES pH 7.4, 150 mM NaCl, 0.01% LMNG buffer and bound proteins were eluted with 2X SDS loading buffer. Samples were run separated using SDS-PAGE (12% gel) followed by Western blotting using HRP coupled M2 anti-FLAG antibody at 1:5,000 dilution.

**HDX-MS analysis.** βarr2 protein samples were prepared in 100 μM as a final concentration in 20 mM HEPES pH 7.4 and 150 mM NaCl. For peptide binding, 500 μM of peptide were added in βarr2 and incubated for 1 hour at room temperature. Hydrogen/deuterium exchange was initiated by mixing 2 μL of protein samples with 28 μL of $D_2O$ buffer (20 mM HEPES pD 7.4, 150 mM NaCl, and 10% Glycerol in $D_2O$) and incubating for 10, 100, 1000, or 10000 seconds on ice. At the indicated time points, the reaction was slowed down by adding 30 μL of ice-cold quench buffer (100 mM $NaH_2PO_4$ pH 2.01). For non-deuterated samples (ND), 2 μL of protein samples were mixed with 28 μL of $H_2O$ buffer (20 mM HEPES pH 7.4, 150 mM NaCl in $H_2O$) and quenched with 30 μL of ice-cold quench buffer. The quenched samples were digested online by passing through an immobilized pepsin column (2.1 x 30 mm) at a flow rate of 100 mL/min with 0.05% formic acid in $H_2O$ at 12℃. Peptide fragments were subsequently collected on a C18 VanGuard trap column (1.7 mm x 30 mm) for desalting with 0.05% formic acid in $H_2O$. Proteins were then separated by ultra-pressure liquid chromatography over an ACQUITY UPLC C18 column (1.7 mm, 1.0 mm x 100 mm) at a flow rate of 40 mL/min with an acetonitrile gradient created by two pumps, which started with 8% B and increased to 85% B over the next 8.5 minutes. Mobile phase A was 0.15% formic acid in $H_2O$, and B was 0.15% formic acid in acetonitrile. To minimize the back-exchange of deuterium to hydrogen, the sample, solvents, trap, and UPLC column were all maintained at pH 2.5 and 0.5℃ during analysis. Mass spectral analyses were performed with a Xevo G2 Qtof equipped with a standard ESI source (Waters, Milford, MA, USA). The mass spectra were acquired in the range of m/z 100–2000 for 12 minutes in positive ion mode. Peptides were identified in non-deuterated samples with ProteinLynx Global Server (PLGS) 2.4 (Waters, Milford, MA, USA). The following parameters were applied: monoisotopic mass, nonspecific for the enzyme while allowing up to 1 missed cleavage, MS/MS ion searches, automatic fragment mass tolerance, and automatic peptide mass tolerance. Searches were performed with the variable methionine oxidation modification, and the peptides were filtered with a peptide score of 6. To process HDX-MS data, the amount of deuterium in each peptide was determined by measuring the centroid of the isotopic distribution using DynamX 2.0 (Waters, Milford, MA, USA). All measurements were performed with three independent experiments, and statistical significance was analyzed by One-way ANOVA test. Back-exchange level was not corrected because all the analysis was comparison of different states.

**Statistical analysis.** For analysis of the time series data, repeated measures ANOVA (rANOVA) was employed at an $\alpha$ level =.01 and the F statistic calculated; time series as a whole were considered to be significant if the F statistic was greater than 1 at the significance level tested. T-test was used to determine the significance between individual time points of the series. When time series data did not meet the threshold of significance by rANOVA, an unpaired or paired (Student's) t-test was employed to assess the significance between time points. GraphPad Prism was used for the statistical analysis.

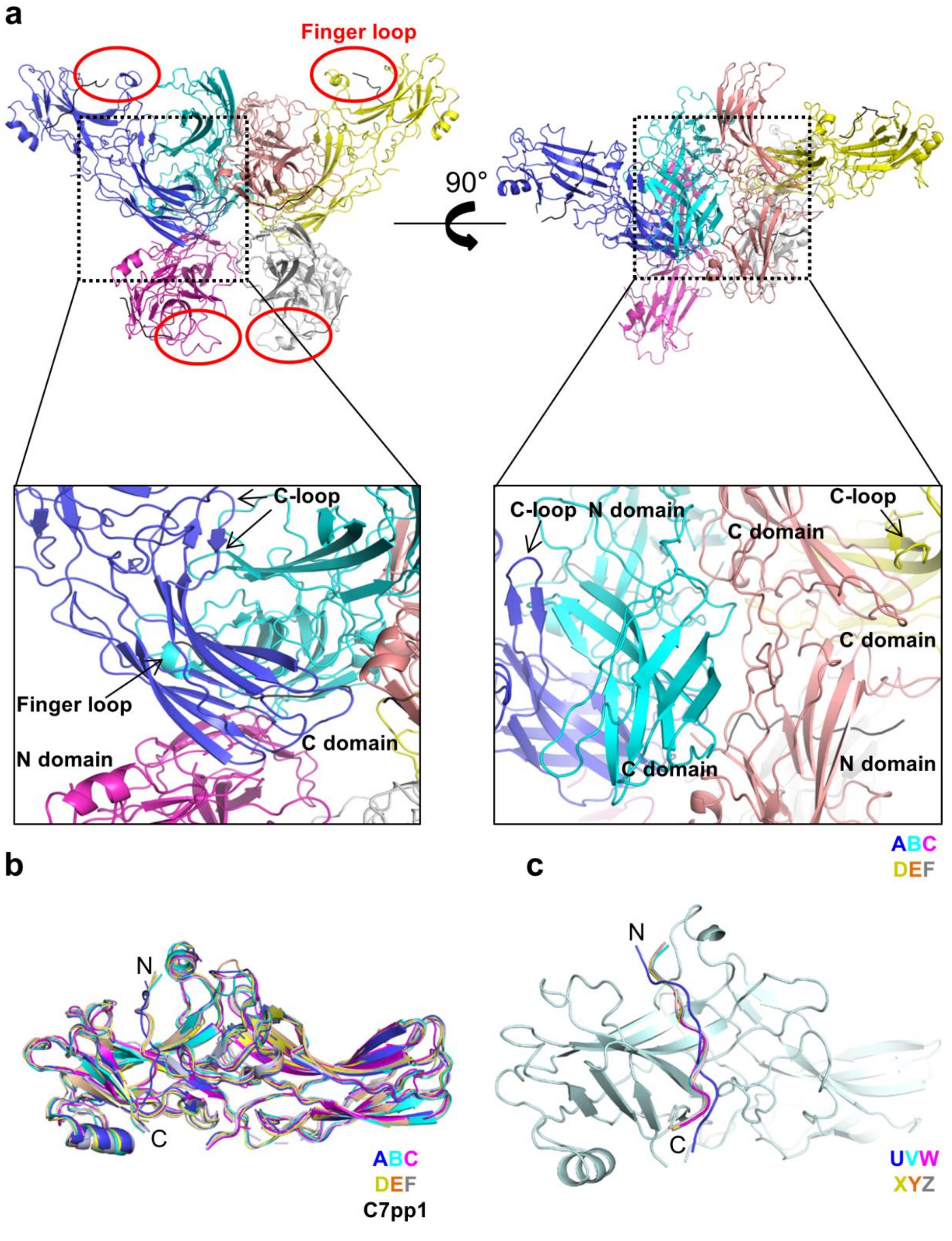


**Supplementary Figure 1. Crystal contacts of the βarr2-C7pp1 complex in asymmetric unit.**

**a**, In $C2_1$ crystallographic lattice, six molecules of the βarr2-C7pp1 complex are contained in an asymmetric unit**.** Each chain is shown with different colors; chain A (light blue), chain B (cyan), chain C (magenta), chain D (olive), chain E (orange), and chain F (grey). Dotted boxes represent magnified crystal contacts between chains. Red circles show solvent exposure of finger loop. Ribbon and surface diagrams of the six molecules rotated by 90° around the indicated axis in the left figure are drawn. The C7pp1 peptides are shown in black lines. **b**, All six chains of the βarr2-C7pp1 complex in asymmetric unit are superimposed. Each chain is shown with different colors; chain A (light blue), chain B (cyan), chain C (magenta), chain D (olive), chain E (orange), and chain F (grey). **c**, The N-domain of each chain is superposed to show different orientations of C7pp1 peptides in each chain. Each C7pp1 peptide represents as different colors; C7pp1 peptide U bound to chain A (light blue), C7pp1 peptide V bound to chain B (cyan), C7pp1 peptide W bound to chain C (magenta), C7pp1 peptide X bound to chain D (olive), C7pp1 peptide Y bound to chain E (orange), and C7pp1 peptide Z bound to chain F (grey).

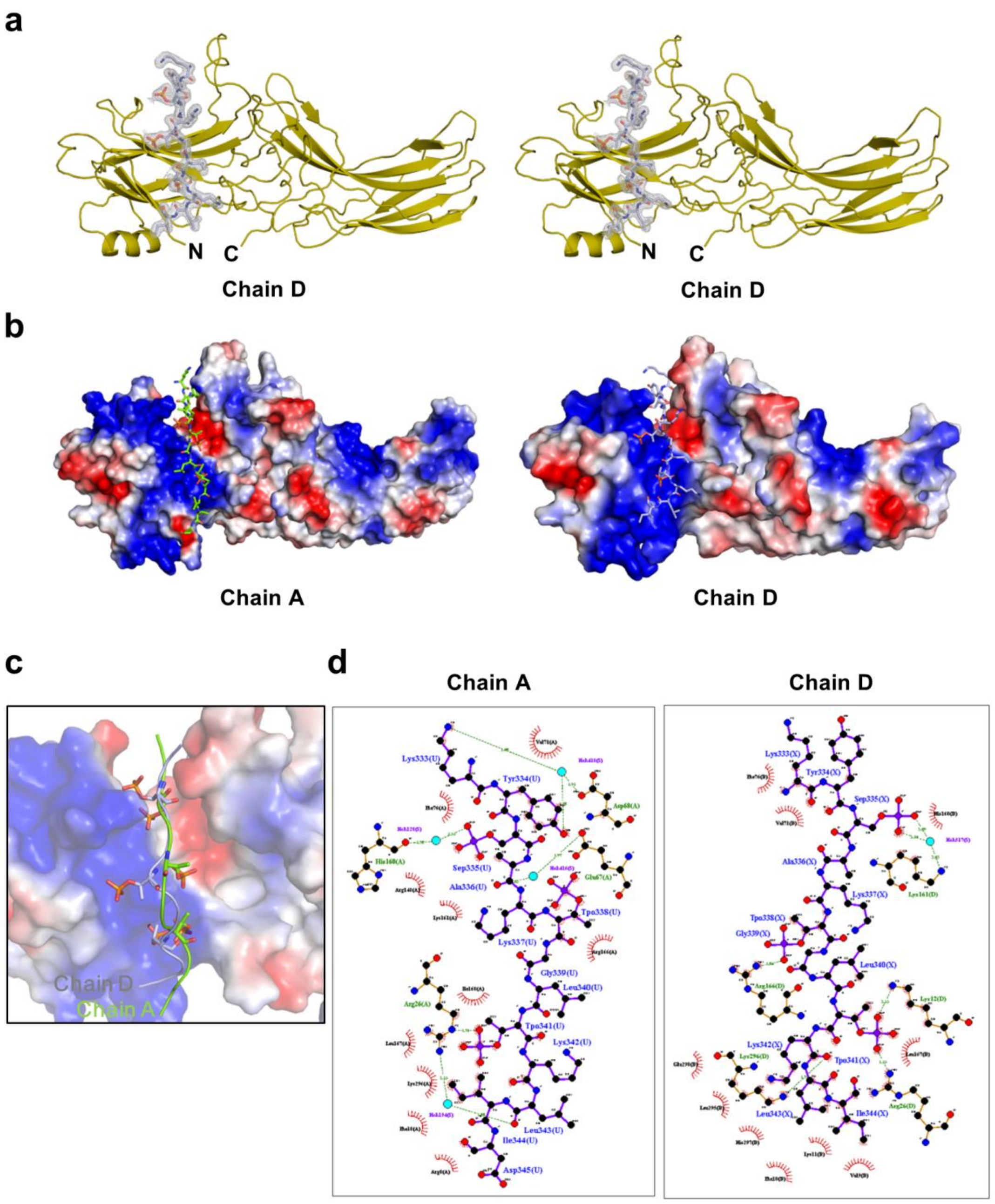


**Supplementary Figure 2. Structural comparison of C7pp1 peptides bound to chain A and chain D. a**, Stereo diagrams of the C7pp1 peptide (chain X) bound to βarr2 (chain D). The βarr2 and C7pp1 peptide are shown as olive and light blue color, respectively. A 2$F_o$ - $F_c$ electron density map (1.0 σ) for C7pp1 is represented. **b**, The electrostatic potential representations for each chain (A and D) of βarr2 are shown. The C7pp1 peptides (chains U and X) bound to βarr2 (chains A and D) are shown as green and light blue sticks, respectively. **c**, Magnified view of both C7pp1 peptides (chains A and D). **d**, Each LIGPLOT diagram[1] represents interactions between βarr2 (chains U and X) and C7pp1 (chains A and D) peptides, respectively.

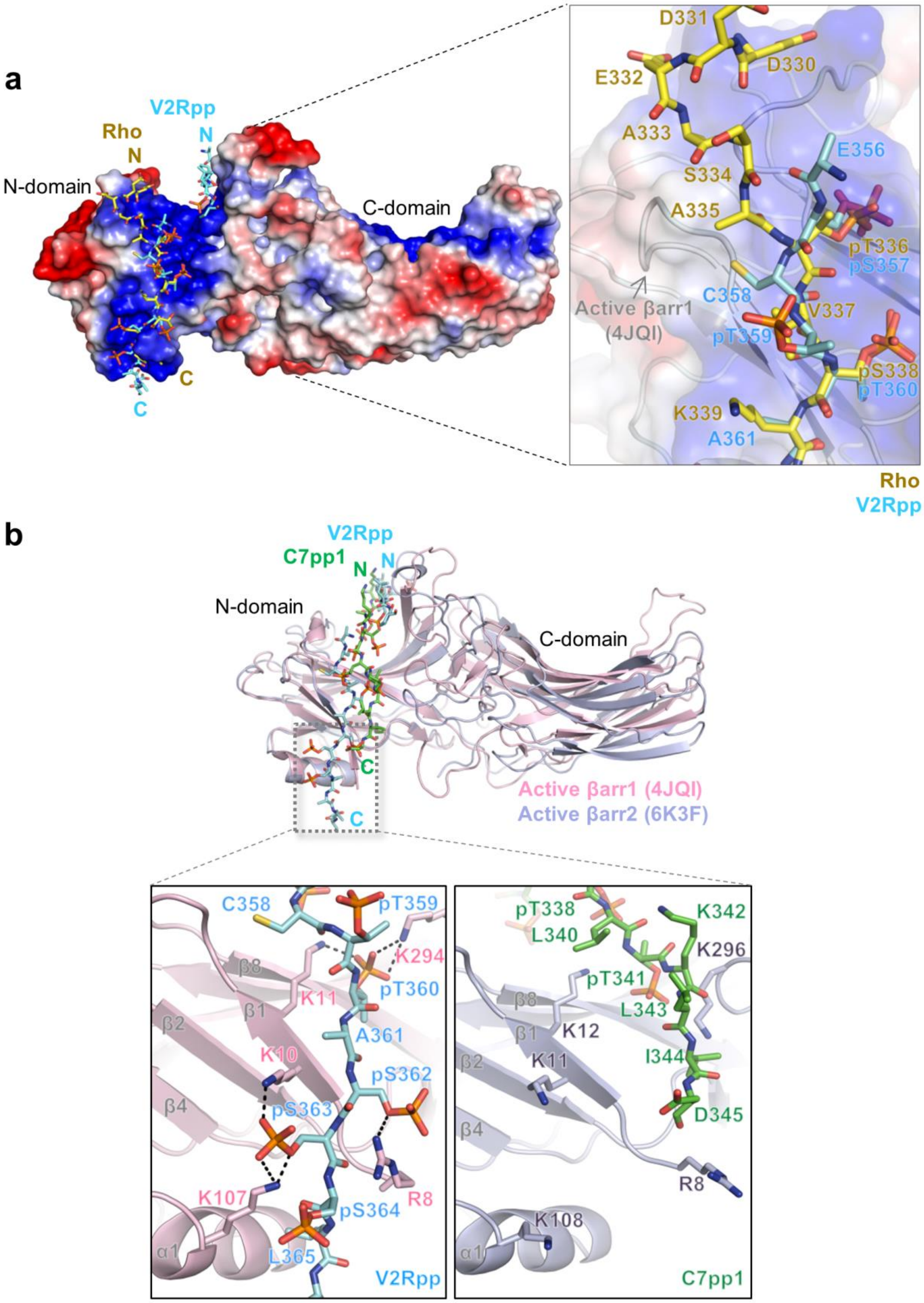


**Supplementary Figure 3. Comparisons of phosphate binding pockets in arrestins. a**, Structural comparisons of phosphate binding sites of $V_2$Rpp and rhodopsin $R_p$-tail (Rho). The

$V_2$Rpp (cyan) and rhodopsin $R_p$-tail (yellow) were superimposed and incorporated into electrostatic potential surface of βarr1 (4JQI). **b**, Structural comparisons of phosphate binding sites of C7pp1 and $V_2$Rpp. Overall structure of the βarr2-C7pp1 complex (6K3F) was represented and $V_2$Rpp was incorporated. The βarr1, βarr2, $V_2$Rpp, and C7pp1 are shown in light pink, light blue, cyan, and green, respectively.

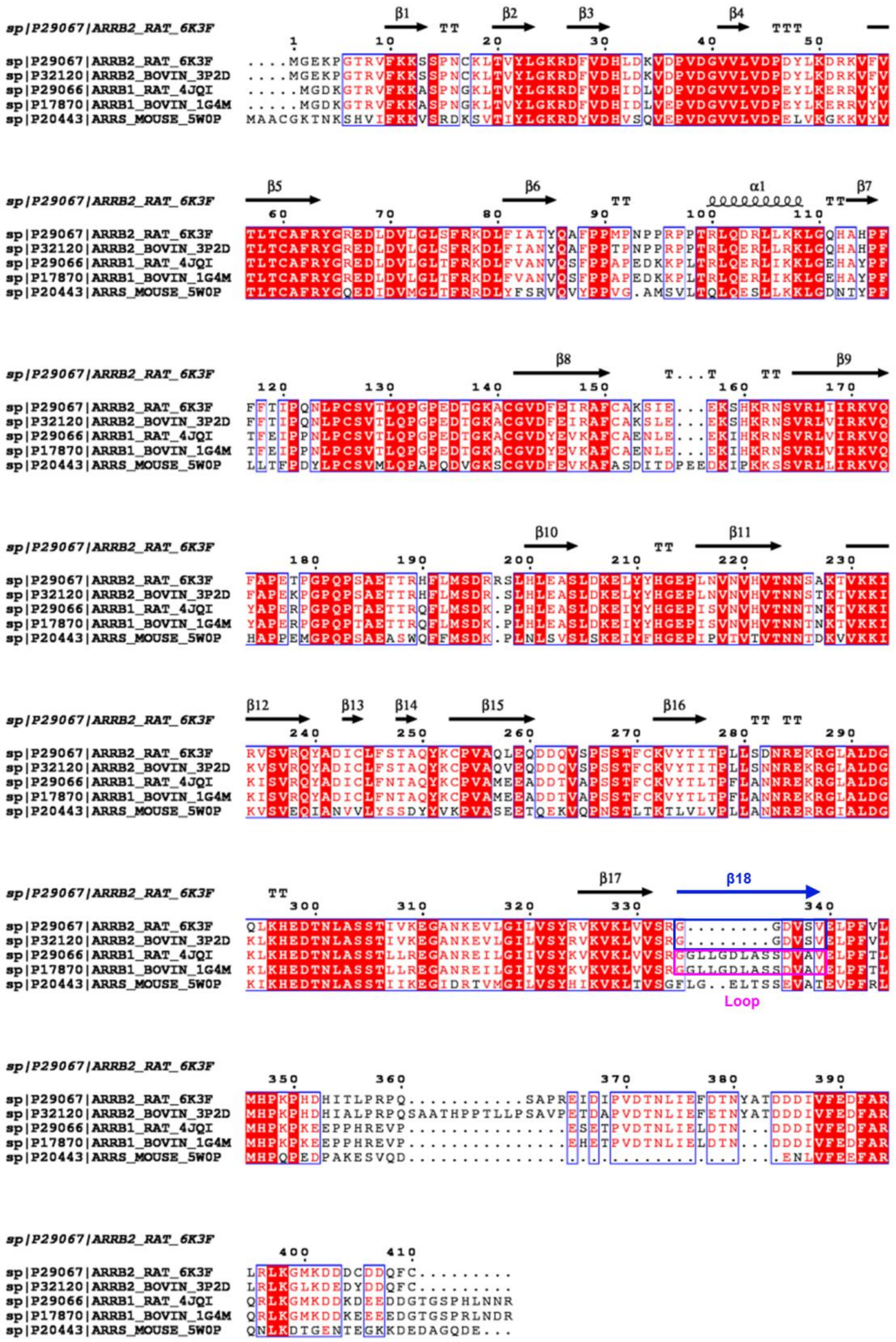


**Supplementary Figure 4. Sequence alignment of βarr2 with other arrestins.** Multi-alignment of *R. norvegicus* βarr2 (UniProtKB/Swiss-Prot accession number P29067, PDB 6K3F) against βarr2 from *Bos taurus* (UniProtKB/Swiss-Prot accession number P32120, PDB 3P2D), βarr1 from *R. norvegicus* (UniProtKB/Swiss-Prot accession number P29066, PDB 4JQI), βarr1 from *B. taurus* (UniProtKB/Swiss-Prot accession number P17870, PDB 1G4M), and S-arrestin from *Mus musculus* (UniProtKB/Swiss-Prot accession number P20443, PDB 5W0P). Secondary structural elements were assigned using PyMOL and every tenth residue is marked with a black dot. Strictly (100%) conserved residues highlighted in red box with white color of characters and semi-conserved residues (80%) are highlighted with red color of characters. Spiral shape and arrows above the sequences denote α-helices and β-strands, respectively. The boxes denoted as blue and pink colors represent additional residues in βarr1 relative to βarr2.

**Supplementary Table 1. Statistics for data collection and refinement.**

| Data set | βarr2 with C7pp1 |
|---|---|
| ***A. Data collection statistics*** | |
| X-ray source | SPring-8 26B |
| X-ray wavelength (Å) | 0.97928 |
| Space group | $C2_1$ |
| *a*, *b*, *c* (Å) | 91.17, 127.91, 206.04 |
| Resolution range (Å) | 50-1.95 |
| Total / unique reflections | 538,906 / 332,324 |
| Completeness (%) | 98.1 (96.3)[a] |
| Average $I/\sigma(I)$ | 70.7 (2.5)[a] |
| $R_{merge}$[b] (%) | 39.0 (150.9)[a] |
| ***B. Model refinement statistics*** | |
| Resolution range (Å) | 50-1.95 |
| $R_{work}$ / $R_{free}$[c] (%) | 18.97 / 19.02 |
| Number / average *B*-factor (Å$^2$) | |
| Protein nonhydrogen atoms | 16,678 / 29.34 |
| Water oxygen atoms | 743 / 28.46 |
| Peptide nonhydrogen atoms | 340/ 28.34 |
| R.m.s. deviations from ideal geometry | |
| Bond lengths (Å) | 0.005 |
| Bond angles (°) | 1.245 |
| Protein-geometry analysis | |
| Ramachandran favored (%) | 80.4 (1669/2077) |
| Ramachandran allowed (%) | 13.9 (288/2077) |
| Ramachandran outliers (%) | 5.7 (120/2077) |

**Footnotes for Table S1**

[a] Values in parentheses refer to the highest resolution shell (1.95-1.98 Å ).

[b] $R_{merge} = \Sigma_{hkl}\Sigma_i \mid I_i(hkl) - \langle I(hkl)\rangle \mid / \Sigma_{hkl}\Sigma_i I_i(hkl)_i$, where $I(hkl)$ is the intensity of reflection *hkl*, $\Sigma_{hkl}$ is the sum over all reflections, and $\Sigma_i$ is the sum over i measurements of reflection *hkl*.

[c] $R = \Sigma_{hkl} \mid |F_{obs}| - |F_{calc}| \mid / \Sigma_{hkl} |F_{obs}|$, where $R_{free}$ was calculated for a randomly chosen 5% of reflections, which were not used for structure refinement and $R_{work}$ was calculated for the remaining.
